# Mutualism reduces the severity of gene disruptions in predictable ways across microbial communities

**DOI:** 10.1101/2023.05.08.539835

**Authors:** Jonathan N. V. Martinson, Jeremy M. Chacón, Brian A. Smith, Alex R. Villarreal, Ryan C. Hunter, William R. Harcombe

## Abstract

Predicting evolution in microbial communities is critical for problems from human health to global nutrient cycling. Understanding how species interactions impact the distribution of fitness effects for a focal population would enhance our ability to predict evolution. Specifically, it would be useful to know if the type of ecological interaction, such as mutualism or competition, changes the average effect of a mutation (i.e., the mean of the distribution of fitness effects). Furthermore, how often does increasing community complexity alter the impact of species interactions on mutant fitness? To address these questions, we created a transposon mutant library in *Salmonella enterica* and measured the fitness of loss of function mutations in 3,550 genes when grown alone versus competitive co-culture or mutualistic co-culture with *Escherichia coli* and *Methylorubrum extorquens.* We found that mutualism reduces the average impact of mutations, while competition had no effect. Additionally, mutant fitness in the 3-species communities can be predicted by averaging the fitness in each 2-species community. Finally, the fitness effects of several knockouts in the mutualistic communities were surprising. We discovered that *S. enterica* is obtaining a different source of carbon and more vitamins and amino acids than we had expected. Our results suggest that species interactions can predictably impact fitness effect distributions, in turn suggesting that evolution may ultimately be predictable in multi-species communities.

## Introduction

Ecological interactions between species shape the evolution of all organisms, yet the impact of these interactions on evolution remains an area of active investigation (1–3). Predicting evolutionary dynamics in microbial communities is critical for everything from understanding how pathogens in the human microbiome will respond to antibiotic treatment (4), to understanding how microbial contributions to global nutrient cycles will be impacted by climate change (5). To understand and predict evolutionary outcomes in microbial communities, it is critical that we gain an understanding of how species interactions impact the average effect of mutations, the specific genes under selection, and how these measurements are impacted as community complexity increases.

One way to assess the impact of species interactions on evolution is to evaluate impacts on the distribution of fitness effects caused by mutations in different conditions (6,7). By measuring the fitness of many mutants one can generate a distribution of fitness effects for a population in a defined environment (6,7). The mean of this distribution is a measure of genetic robustness and indicates the average effect of a mutation in a population in the defined environment. One can evaluate how environments alter the average effect of mutations by comparing the mean of the distribution of fitness effects measured in two different environments. One approach that can be used to measure the distribution of fitness effects is known as TnSeq (8). In this method, transposon mutant libraries are generated that contain thousands of mutants, each harboring a transposon inserted in a distinct location in the genome (9,10). Through sequencing the transposon library before and after growth under defined conditions, one can determine the change in mutant frequency and thereby the relative fitness of each insertion. While TnSeq studies typically focus just on outliers to identify essential genes in different environments (9), the field is beginning to use the technique to study the impacts on the entire distribution of fitness effects for a given species (7,11–13). Indeed, the average effect of mutations on a given strain is often observed to change in different environments (14).

Recent studies have shown that species interactions can change the fitness effects of mutations in a variety of ways, but it remains unclear how well pair-wise effects can be predicted. For instance, a TnSeq study with *E. coli* found that knockouts of several vitamin and nucleotide biosynthesis genes were deadly in monoculture, but viable in the presence of a phototrophic bacterium (12). Morin *et al.* tested the same *E. coli* library with different species of bacteria and fungi, and found that interactions with other species can result in either alleviation or exacerbation of fitness costs caused by transposon insertion (13,15). These two studies demonstrate that fitness effects of species interactions can be idiosyncratic, however there may be some generalities based on how species interact. Specifically, competition may tend to render gene disruptions more deleterious by further penalizing slow growth, while mutualism may tend to buffer the effect of gene loss through constraining growth rates to match those of a partner. To evaluate this possibility, data are needed on mutational effects in a system in which species identity can be held constant, while species interactions can be changed.

Another open question is whether the impact of mutations in complex communities can be predicted from the impact of mutations in simpler systems. Morin *et al.* found that only 28% of the fitness effects in a 4-species community could be predicted from additive combinations of effects in pairwise interactions (15). This work suggests that higher order interactions (i.e., emergent properties) are prevalent and that the impact of mutations in communities cannot be well predicted from pairwise interactions. However, this contrasts with many studies suggesting that higher order interactions are relatively rare in microbial communities (16,17). The predictability of mutational impacts in a community setting remains unclear.

Here we use a randomly barcoded transposon-mutant (RB-TnSeq) library of *Salmonella enterica* (abbreviated as ‘S’) to investigate how species interactions impact the average effect of mutations, and the effect of disrupting specific genes. Changes in the nutrients provided in the media were used to switch *S. enterica* between mutualism and competition with *Escherichia coli* and *Methylorubrum extorquens* (abbreviated as ‘E’ and ‘M’, respectively). Our strain of *S. enterica* secretes methionine (18). In lactose minimal medium, this *S. enterica* strain forms an obligate mutualism with an *E. coli* methionine auxotroph by providing methionine in exchange for carbon byproducts (Fig. 1A) (19). The *S. enterica* can also form an obligate mutualism with *M. extorquens* in galactose minimal medium providing carbon byproducts in exchange for nitrogen when methylamine is the only nitrogen source (20). Thus, all three species form an obligate mutualism in lactose minimal medium with methylamine (Fig. 1A). However, if succinate, methionine, and ammonium are each provided in the medium, all three strains can grow independently and act as competitors of each other (Fig. 1A). We grew the *S. enterica* RB-TnSeq library alone, with *E. coli*, with *M. extorquens* and with both species in either mutualism or competition and sequenced the transposon barcodes in a process called BarSeq (10). Monocultures were grown in galactose minimal medium for comparison against mutualistic growth, and succinate minimal medium to compare against competitive growth. We compared the effect of co-culture relative to its paired monoculture to determine whether the distribution of fitness effects varies in competition versus mutualism.

**Figure 1.**
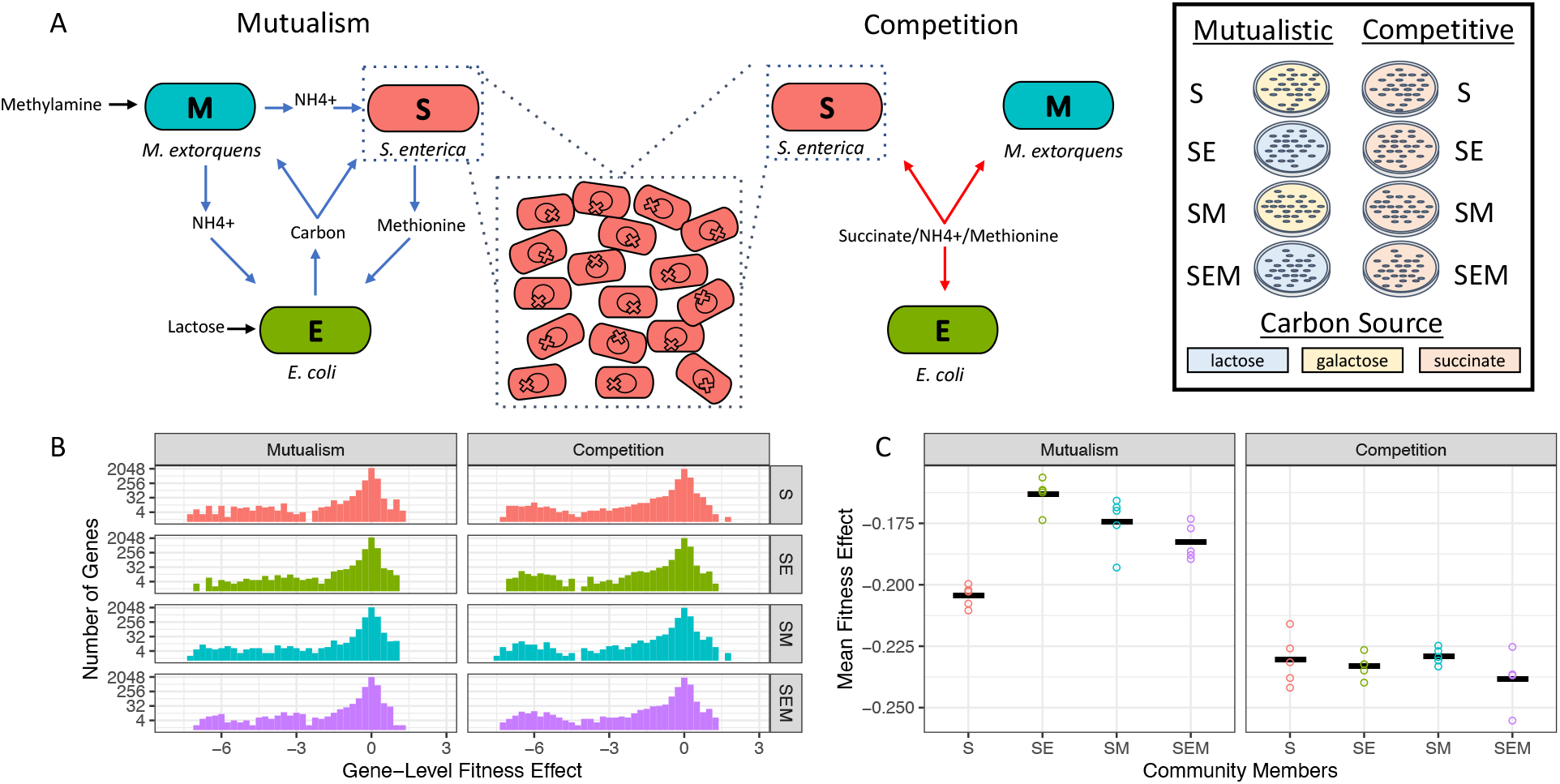
Distributions of fitness effects during mutualism and competition. A) Interactions between *S. enterica* (S), *E. coli* (E), and *M. extorquens* (M) can be switched from mutualism (left) to competition (right) by changing the media. Central dotted inset – *S. enterica* population is composed of a transposon insertion (gene knockout) library. The circle within each cell represents a DNA chromosome – each X represents a transposon insertion site (knockout). The library was tested in monoculture, 2-species co-culture, and 3-species co-culture (rightmost box). Plate color indicates carbon source – yellow = galactose; blue = lactose; orange = succinate. B) Histograms of gene-level fitness effects. Each gene’s fitness effect was averaged over the 5 replicates within the treatment. Fitness effect is the normalized fitness as described in the Materials, such that a fitness effect = 0 indicates a neutral knockout. C) Means of the fitness effects over all genes within a replicate. Each point is the mean of one replicate’s fitness effects. The horizontal line is the mean of these means. The mean fitness effect of knockouts in S monoculture was significantly (p = 2e-10) negative. There was a significant increase in the mean fitness effect when S depended on E (p = 2e-7) or M (p = 1e-5). Dependence on both E and M did not additively increase mean fitness effect, and instead lowered the mean below that for dependence on either species (p = 1.8e-6). The mean fitness effect of knockouts in competitive treatments was significantly below zero (intercept p < 2e-16) and the community members had no effect (smallest p = 0.144).

We found that species interactions changed both the average effect of disrupting a gene as well as which genes were under selection. When *S. enterica* engaged in mutualism it was less affected by gene deletions than in monoculture, while in the competitive environment species interactions had no significant impact on the average fitness effect of gene disruption. The BarSeq data highlighted some expected interaction-specific selection such as the increased selection on genes associated with nitrogen uptake when *S. enterica* was reliant on nitrogen from *M. extorquens* (12). The BarSeq data also illuminated some unanticipated selection, such as changes in the importance of vitamin biosynthesis in mutualism. Encouragingly, our data suggest that selection in 3-species communities can be well predicted from fitness in pairwise associations.

## Materials and Methods

### Bacterial strains

Strains are listed in table S3. *Salmonella enterica* LT2 (WH102), and *Methylorubrum extorquens* AM1 have been described in previous studies (19,21). Briefly, the *S. enterica* contains mutations in *metA* and *metJ* causing it to secrete methionine (18), and the *M. extorquens* has a deletion of *hprA* making it unable to assimilate carbon from methylamine. The *Escherichia coli* strain was generated using a P1 transduction method (22,23) to move the *metB* deletion from the Keio clone JW3910 into *Escherichia coli* MG1655. The resistance cassette was removed with flippase (22,23).

Single gene knockouts in *S. enterica* were constructed using P22 HT *int* transduction to move knockouts from the BEI Resources *S. enterica* 14028s knockout library. BEI plate IDs for the strains used were: Δ*aceA* –SGD_156/157_Cm, NR-42890 well A02; Δ*panC* – SGD_051/052_Cm, NR-42877, well D10; Δ*ilvA* – SGD_156/157_Cm, NR-42890 well F08. Transductions were carried out in a similar fashion to the P1 transductions described above, however, antibiotic resistance cassettes were left intact.

### Media

Routine culturing of *E. coli* (E) and *S. enterica* (S) was carried out on Lysogeny broth (LB), while *M. extorquens* (M) was cultured on Nutrient broth. The BarSeq experiments were carried out on Hypho minimal medium (24). Modified Hypho minimal medium was prepared as shown in Supplemental Table 1. Each component was sterilized before mixing. The carbon source of the mutualistic communities was either 5.56 mM galactose (for S and SM) or 2.78 mM lactose (for SE and SEM). Competitive communities were provided 8.33 mM succinate as the carbon source.

Mutualistic communities without *M. extorquens* (S and SE) and the competitive communities were provided (NH_4_)_2_SO_4_ (3.7 mM) as the nitrogen source. Mutualistic communities with *M. extorquens* (SM and SEM) were provided Na_2_SO_4_ (3.78 mM) and 1.16 mM methylamine. The competitive medium was supplemented with 0.05 mM methionine to allow for unrestricted *E coli* growth. All media was supplemented with 1.2 µM ZnSO_4_, 1 µM MnCl_2_, 18 µM FeSO_4_, 2 µM (NH_4_)_6_Mo_7_O_24_, 1 µM CuSO_4_, 2 mM CoCl_2_, 0.33 µM Na_2_WO_4_, and 20 µM CaCl_2_.

### RB-TnSeq library construction for *Salmonella enterica*

A randomly barcoded transposon library (RB-TnSeq) was generated in *S. enterica* WH102 using the conjugation method described in Wetmore *et al*. (10). Briefly, the donor *E. coli* strain APA752 containing the suicide transposon-plasmid, pKMW3, was conjugated with WH102 at a 1:1 ratio on LB supplemented with 300µM diaminopimelic acid for 24 hours at 37°C. Roughly 300,000 transconjugant colonies were scraped and resuspended in saline (0.9%). The cells were then diluted to OD600 0.25 in LB with kanamycin (50µg/mL) and grown to an OD600 of 0.97 prior to being frozen in 10% glycerol at -80°C. The library was sequenced on a NovaSeq S1 2x150bp flow cell at the University of Minnesota Genomics Center and analyzed using the FEBA pipeline (10) against the *S. enterica* LT2 genome (GenBank accession: AE006468.2).

### BarSeq Experimental Setup

A 1 mL RB-TnSeq library aliquot was thawed and inoculated into 25 mL LB with 50µg/mL kanamycin and incubated at 37°C until it reached mid-log phase (OD600 0.64). *E. coli* (MG1655 Δ*metB*) was cultured overnight and then diluted to mid-log in LB 37°C. *M. exotoruquens* Δ*hprA* AM1 was cultured for two days in nutrient broth at 30°C. All species were washed and adjusted to OD600=0.2 (for *S. enterica* and *E. coli*) or 0.4 (for *M. extorquens)*. 25 µL of each species (∼1E6 cells) was plated onto minimal media (5 replicates/treatment) then incubated at 30°C.

To harvest cells from the experimental plates, 4 mL saline was pipetted into each replicate plate and colonies were scraped with a cell spreader. A sample of the cell slurry was plated for CFU quantification on modified Hypho agar (Supplemental Table 2). The remaining cell slurry was pelleted by centrifugation and frozen at -20° C.

DNA was extracted from pelleted cells using the Qiagen DNeasy Blood and Tissue kit. The resultant DNA was quantified with the Quant-iT™ PicoGreen™ dsDNA Assay kit. We performed the BarSeq98 PCR method previously described (10) with NEB Q5 polymerase and 200ng of genomic DNA. In total, 55 samples with unique sequencing index barcodes were sequenced on a single lane on an Illumina NextSeq P2 1x100-bp run that generated 315 million reads.

### Data filtering

Data filtering followed much of the same procedure described previously (10). First, we removed barcodes which were not in chromosomal genes. Second, we removed barcodes that fell in the first, or last, 10% of a gene. Third, we removed barcodes which had three or fewer reads in any Time 0 (T0) sample. All samples had a median of more than 50 reads per gene. Fourth, we removed genes from the analysis if any T0 sample had fewer than 30 reads total in a gene, or fewer than 15 reads in the first or last half of the gene. This left 105,000 barcodes across 3,550 genes. Note that one sample failed to pass quality check scores (competitive SEM replicate 1) and was removed from the analysis.

### Barcode-level fitness calculation

We calculated barcode-level fitness using a hybrid of the approaches described previously (10,25). First, a pseudocount of 0.1 was added to each barcode count to avoid taking log of zeros. Second, we divided each replicate’s read counts by the average number of barcode reads in five reference genes which we expected to have no fitness effect when knocked-out; this was done per-sample (the reference genes were STM0604, STM1237, STM0329, STM2774, and STM0333). To put the normalized counts back onto the original count scale, we multiplied the within sample normalized counts by the average number of barcode reads in the five reference genes over all samples. Finally, barcode-level fitness was calculated by log2(normalized counts) – log2(normalized counts of T0 sample). There were five T0 samples. Each sample within each treatment was designated as replicate 1-5, and each used a different T0 sample for the fitness calculation. Barcode-level fitness variance was calculated as previously performed (10).

### Gene-level fitness calculation

Fitness for each gene within each replicate was calculated as a weighted average of barcode-level fitness calculations. Specifically, fitness was weighted by 1 / barcode-level variance. We set the maximum weight a barcode could receive at 20 reads (10). Gene-level fitness was then Σ (barcode fitnesses X barcode weights) / Σ (barcode weights).

Finally, we corrected for chromosome position. Following (10), within each sample, we calculated a rolling median of gene-level fitness along the chromosome with a window size of 251 genes. This rolling median was subtracted from each gene’s fitness to obtain the final gene-level fitness value used in downstream analyses.

### Statistics

To compare the mean fitness of treatments, we first found the mean fitness within a replicate by taking the arithmetic average of the fitness of each gene disruption. These means were the response variable in a linear regression including the presence of *E. coli*, of *M. extorquens,* and the interaction term as explanatory variables. Because these are binary variables, the coefficients can be interpreted directly as the change in mean fitness due to the presence of *E. coli* or *M. extorquens*, or (for the interaction term) the change from additivity. We ran separate tests for mutualism and competition.

To compare gene-level fitness values across species addition treatments we used the same approach as above. To correct for multiple comparisons, we controlled the false-discovery rate by adjusting the p-values across genes using Benjamini-Hochberg (BH) corrections and designated adjusted p < 0.05 as significant.

To predict gene fitness in the three-species communities, different methods were tested. The sum, largest, or average effects of E and M terms for each gene were added to the monoculture fitness for each gene. Predictions for all genes were correlated with the observed ESM fitness value and the R^2^ was examined. To compare prediction metrics a bootstrap analysis with 2001 iterations (with resampling) was performed. For each iteration, prediction values and ESM values were resampled, R^2^ values were calculated, and the frequency of times R^2^ values differed for each prediction was calculated. This process was repeated independently for both competition and mutualism.

In the overrepresentation analysis (ORA) we filtered for significant terms (BH adjusted p < 0.05) and classified gene effects as either positive or negative. ORA was then performed separately on the genes classified as positive and negative using the ‘enrichKEGG’ function from the R package, ‘ClusterProfiler’ (version 4.7.1, downloaded 3/1/23) (26) with a universe size of 3550.

### Growth measurements

Growth curves in liquid were performed on overnight cultures grown in minimal medium. Cells were washed 3 times in saline, adjusted to an OD600 of 0.2, and diluted 1:100 in the relevant minimal medium. Cells were incubated at 30°C in a Tecan Infinite Pro200 and shaken at 432 rotations per minute. Growth curve parameters were obtained by fitting a log-linear model to the OD600.

Growth measurements for communities on solid agar were performed by growing bacteria as in the ‘BarSeq Experimental Setup’ except that wildtype *S. enterica* was used in place of the library. Plates were scanned (600 dpi) once per hour on an Epson Perfection V600 scanner at 30°C. Lawn density was calculated by converting to gray scale, performing a Gaussian blur, and measuring the mean gray value for each plate using a custom script written with assistance from GPT-4 (27).

### Spent media preparation

Spent media was prepared by inoculating hypho minimal media with a 1:100 dilution of washed cells grown overnight. *M. extrorquens* spent media was prepared in SM competitive broth (Supplemental Table 1). *E. coli* spent media was prepared in hypho with lactose and methionine. *E. coli* were grown to mid-log phase (OD600 ∼0.2) or stationary phase (OD600 ∼0.4). Spent media was supplemented with 1% galactose, 10% P-solution, and 10% S solution. *S. enterica* spent media was made in either galactose or succinate + methionine hypho. Spent media from mid-log phase was then mixed at a 1:1 ratio with fresh galactose hypho. All spent media was centrifuged and then filter sterilized (0.2 µm).

### Vitamin and amino acid gradients

Galactose or succinate + methionine hypho were supplemented with either calcium pantothenate (0.5µM to 5e-7µM; Fisher Scientific) or DL-isoleucine (1mM to 1e-6 mM; Alfa Aesar). The Δ*panC* and Δ*ilvA* were grown overnight, washed and adjusted to OD600=0.2, and diluted 1:100 before growth was measured.

### High performance liquid chromatography (HPLC) analysis

Quantification of organic acids in spent media was completed using a Dionex UltiMate 3000 RS HPLC system equipped with an Acclaim organic acid (OA) 5μm 120A° 4.0 x 250 mm column and accompanying guard column. Analyte separation was achieved using a 32-minute isocratic run method consisting of an 8-minute equilibration step followed by a 24-minute static flow step with 100mM NaSO_4_ (pH adjusted to pH 2.6 using CH_3_SO_3_H) at a rate of 1mL/min. The RS column oven compartment temperature was maintained at 30°C and the RS diode array was configured to collect UV readings at a wavelength of 210nm with default frequency. All standards and samples were filtered through 0.22μm polyethersulfone (PES) polymer filter membranes prior to injection (6μL per sample). Chromeleon software (v.7.0) was used to configure all run sequence settings as well as view chromatogram data. Analyte peaks were identified, gated, and measured via the integrated Cobra Wizard prior to data export and processing.

### Scripts and Data Availability

Statistics and figure generation were performed using R 4.2.1 and Python 3.9 using custom scripts available at https://github.com/JonMartinson/ecology_DFE. Raw transposon sequencing data are available at SRA #########.

## Results

The overall goal of our study was to determine how ecological interactions impact the distribution of fitness effects of mutation. The BarSeq experiment resulted in 3,550 *S. enterica* gene disruptions which passed quality control (see Methods) and therefore entered downstream analyses. The *S. enterica* populations from which the BarSeq data were obtained experienced a similar number of generations between treatments within each ecological category of mutualism or competition (Supp. Fig. 1), which allowed us to make direct comparisons within a category but not between. In the mutualism treatment all co-cultures grew more slowly than the monoculture (Supp. Fig. 2). *S. enterica* had slightly but significantly lower yield in competitive co-culture than monoculture consistent with weak competition (t-test, p<0.05, Supp. Fig. 1).

### Distribution of Fitness Effects

As expected, most gene disruptions were neutral (relative fitness near zero), with a longer tail towards knockouts causing low fitness than high fitness in all treatments (Fig. 1B). To specifically evaluate the impact of species interactions on the average effect of mutations we performed linear regression analysis. Our model had a factorial treatment structure with the terms ‘presence of E’ and ‘presence of M’ denoting the species composition of each community. The adjusted R^2^ value for the mutualistic community was 0.80 (p < 1e-5, F = 26.43), while the competitive community model’s adjusted R^2^ was 0.01 (p > 0.05, F = 1.09). These results suggest that species composition predicts the mean fitness effect of mutation in mutualistic communities, but not in competitive communities. Full results of both regressions are shown in Supp. Tables 4 and 5.

In all communities (both mutualistic and competitive), the average fitness of mutants was below zero, which is consistent with mutations generally being deleterious. In the mutualistic environment, the mean fitness effect of mutations was less deleterious when *S. enterica* depended on *E. coli* (linear regression model, β = 0.04, p = 2e-7) or *M. extorquens* (linear regression model, β = 0.03, p = 1e-5) compared to when *S. enterica* was grown by itself (Fig. 1C, left). In the competitive treatments species composition did not have a significant impact on mean fitness.

We next investigated what drives the average effect of mutations to be less deleterious when *S. enterica* is engaged in mutualism. Fitness of knockouts in monoculture and mutualistic co-culture were generally similar (R^2^ = 0.87 for S vs. SE, 0.85 for S vs. SM, Fig. 2 AB). However, some knockouts had different fitness in co-culture than in monoculture. In mutualism, when fitness differed from monoculture, it was more likely to increase when grown with either *E. coli* (binomial test, p = 0.002, Fig. 2C) or *M. extorquens* (binomial test, p < 0.001, Fig. 2C). There was a substantial degree of overlap in gene disruptions that were rescued by both *E. coli* and *M. extorquens* in mutualism (Fig. 2C). In contrast, competition had a significant effect in the opposite direction when *S. enterica* was grown with *E. coli* (binomial test, p = 0.005, Fig. 2F) and no significant changes in fitness with *M. extorquens*.

**Figure 2:**
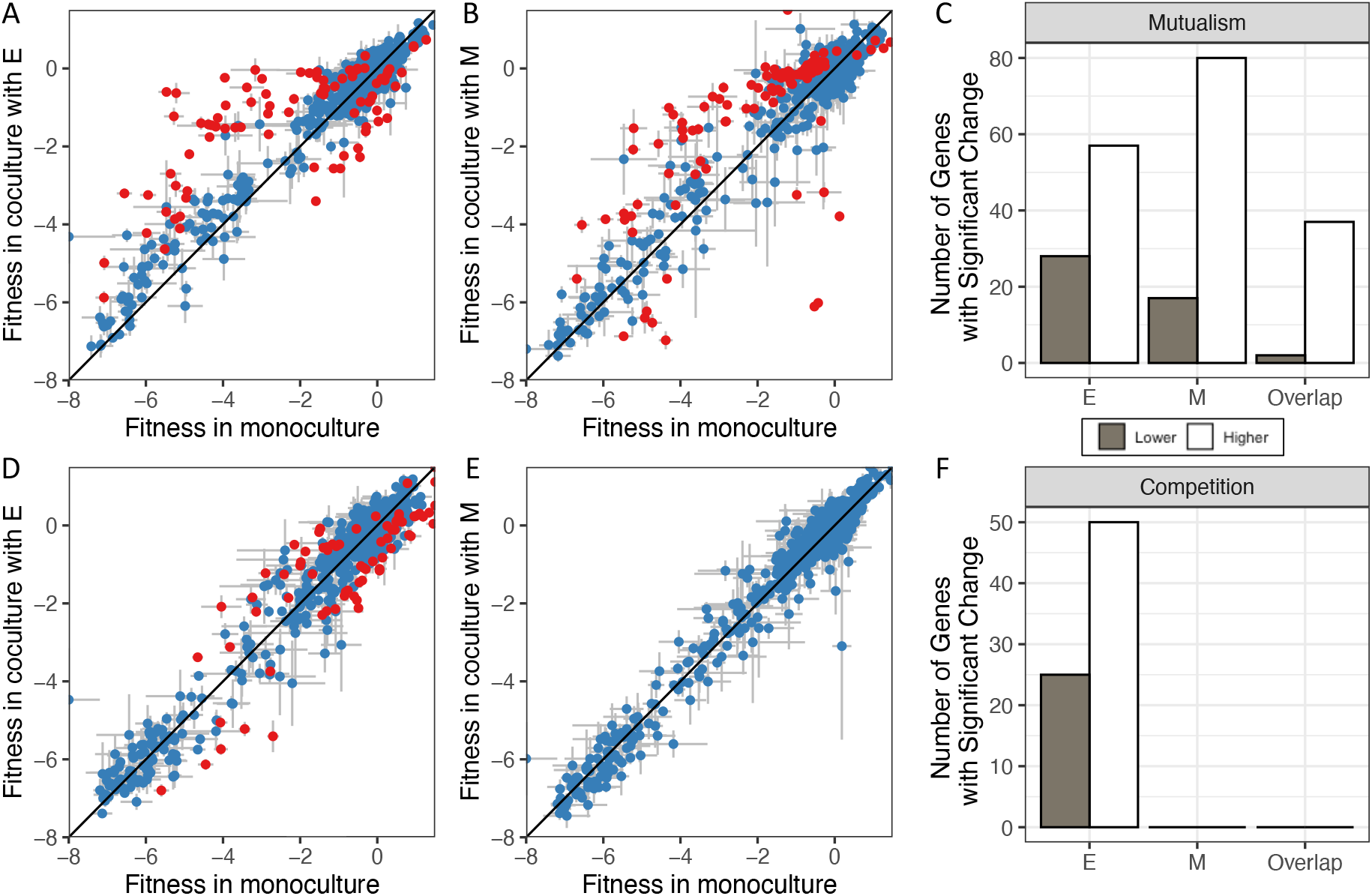
Mutualism tends to rescue mutants while competition slightly decreases mutant fitness. The mean fitness effect of each gene knockout in monoculture vs. co-culture with *E. coli* (A,D) or *M. extorquens* (B,E). The top row shows mutualistic co-cultures while the bottom row shows competitive co-cultures. Each point represents the fitness of one gene knockout. The error bars are the standard error of the mean over 5 replicates. Red dots indicate the co-culture fitness value differed significantly from the monoculture fitness value. (C,F) The number of genes that that had significant change in each co-culture, and the overlap in these genes. Solid bars represent the number of genes that had significantly lower fitness in co-culture, while open bars represent genes with significantly higher fitness in co-culture than monoculture.

### Predicting effects of increasing community complexity

We then tested whether fitness in the 3-species community could be predicted from the fitness in each 2-species community. We specifically evaluated whether fitness in the 3-species communities could be predicted from: i) the additive effect on fitness of a knockout in each 2-species co-culture ii) the strongest effect on fitness in either 2-species co-culture, or iii) the average effect on fitness in each 2-species co-culture. All three prediction methods had high R^2^ (median > 0.8); however, the R^2^ was highest for predictions based on average fitness effects in both mutualism (Fig. 3A left) and competition (Fig. 3A right). The differences were slight particularly for competition, but a bootstrap analysis indicates a significant difference between predictive metrics for both mutualism and competition (bootstrap p < 0.05).

**Figure 3.**
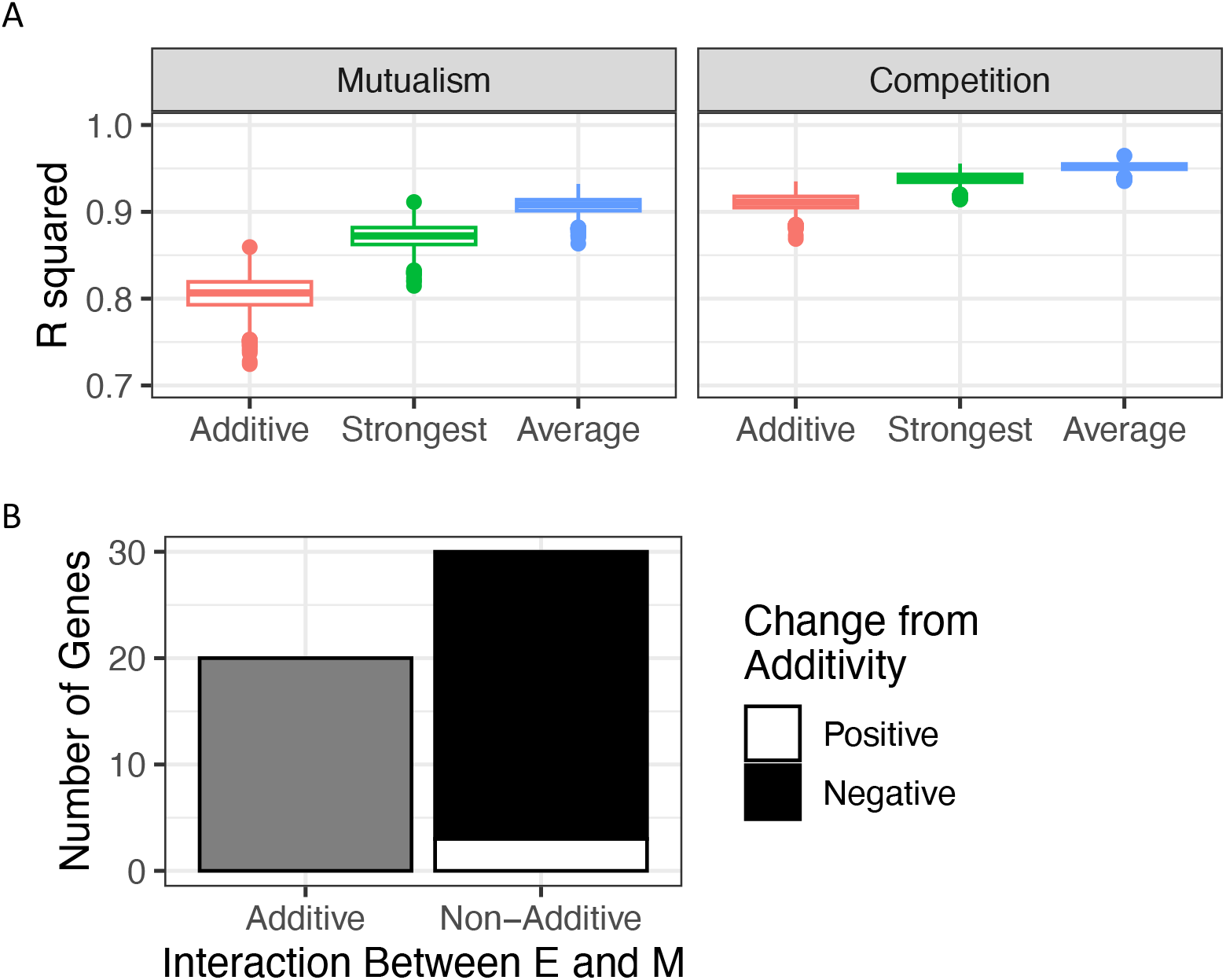
Predictability of fitness effects by community composition. a) Distributions of 2001 bootstrapped R-squared values for Additive, Strongest, and Average models in predicting gene fitness in three-species communities under mutualistic (left) and competitive (right) conditions. b) The number of genes for which both E and M caused a significant main effect (in mutualism), scored by whether the interaction term was also significant (i.e. non-additive). The significant interaction term data are further classified into whether the fitness in SEM is higher (positive – blue) or lower (negative – red) than expected from additivity.

The fitness effects of knockouts in each co-culture tended to be sub-additive in the 3-species mutualism. When both *E. coli* and *M. extorquens* influenced the fitness of a knockout in their respective 2-species co-cultures, 30 out of the 50 knockouts had non-additive effects of adding both species (Fig. 3B). Of the 30 knockouts with non-additive effects, 27 exhibited sub-additivity: the typical result was that each co-culture species ameliorated the fitness cost of a knockout in comparison to monoculture, but the combined effect of the species in the 3-species co-culture was less than their sum.

### System-specific inference

In addition to investigating how species interactions impacted the distribution of fitness effects, we also sought to investigate how species interactions altered the specific genes that contribute to fitness. We were particularly interested in understanding the mechanisms of interaction within the mutualism so we performed overrepresentation analysis (ORA) to identify pathways that become more and less important as a result of species interactions. We determined the effect of species composition on the fitness of each gene for each ecological treatment using multiple linear regression (Supp. Fig. 3). In the presence of *M. extorquens*, ORA identified the nitrogen metabolism pathways as overrepresented, as expected. Surprisingly, we found that several vitamin and amino acid biosynthesis pathways became less important in the presence of mutualistic partners suggesting the possibility of additional cross-fed metabolites (Fig. 4). Based on this analysis we further investigated the fitness effects of specific genes in pathways whose importance changed as a result of species interactions.

**Figure 4.**
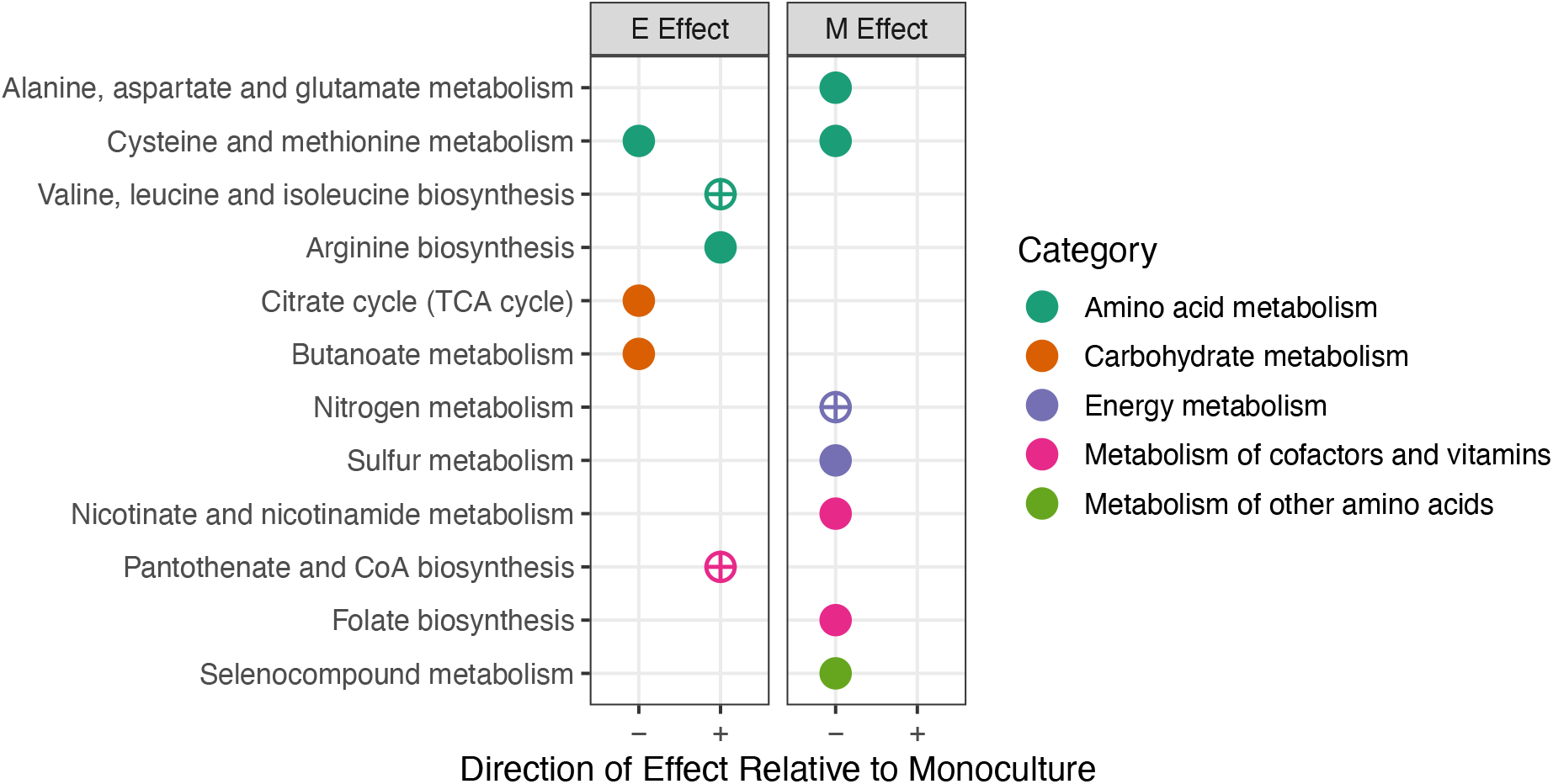
Overrepresentation Analysis of Genes with Significant Fitness Effects. Dotplot of KEGG pathways significantly overrepresented in the set of genes that were significant as an effect of the presence of E or M (p < 0.05, BH multiple comparisons). Pathways overrepresented are indicated by circles. The plus/minus on the x-axis indicate the direction (relative to monoculture) of the fitness effect on the pathway. Empty circles indicate pathways that we investigated further.

### Nitrogen metabolism

As expected, genes associated with nitrogen metabolism were critical in mutualistic co-culture with *M. extorquens. S enterica* depends on NH_4_^+^ from *M. extorquens* when the two species are engaged in mutualism. Genes involved in nitrogen stress, *lrp* and *glnK*, had a significantly stronger negative fitness effect in mutualistic co-culture with *M. extorquens* than in monoculture (Fig. 5). Similarly, an ammonium transporter (*amtB*) and genes involved in glutamate biosynthesis under low ammonium concentrations (*gltB* and *gltD*), had fitness effects that were neutral in monoculture but highly negative in mutualistic co-cultures with *M. extorquens*. These genes had no significant fitness effects in competitive interactions (Fig. 5).

**Figure 5.**
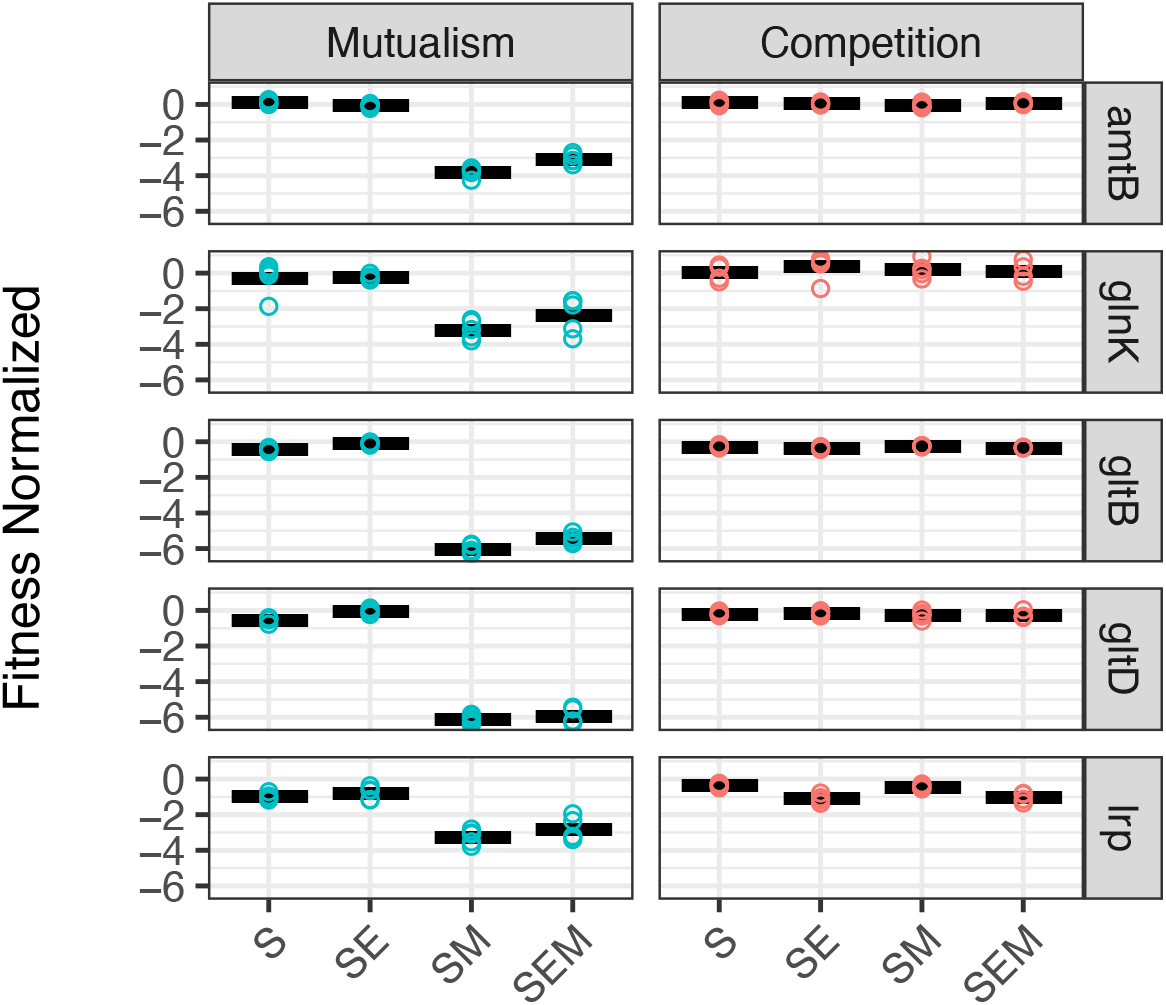
Mutualism with *M. extorquens* results in strong negative fitness effects in nitrogen metabolism genes. Normalized fitness scores for nitrogen metabolism genes in mutualistic (red) and competitive (blue) conditions. Each dot represents the average fitness in a replicate, and the horizontal line represents the mean fitness for each gene across replicates.

### Carbon metabolism

We next investigated the fitness effects of altering carbon metabolism when *S. enterica* was reliant on *E. coli*. Metabolic modeling and previous work led us to expect that *S. enterica* obtains a combination of acetate and galactose from *E. coli*. In contrast to these expectations, we found that transposons that knocked out genes critical for acetate or galactose consumption had little impact on the fitness of *S. enterica* when it was cross-feeding from *E. coli* (Fig. 6 A).

**Figure 6.**
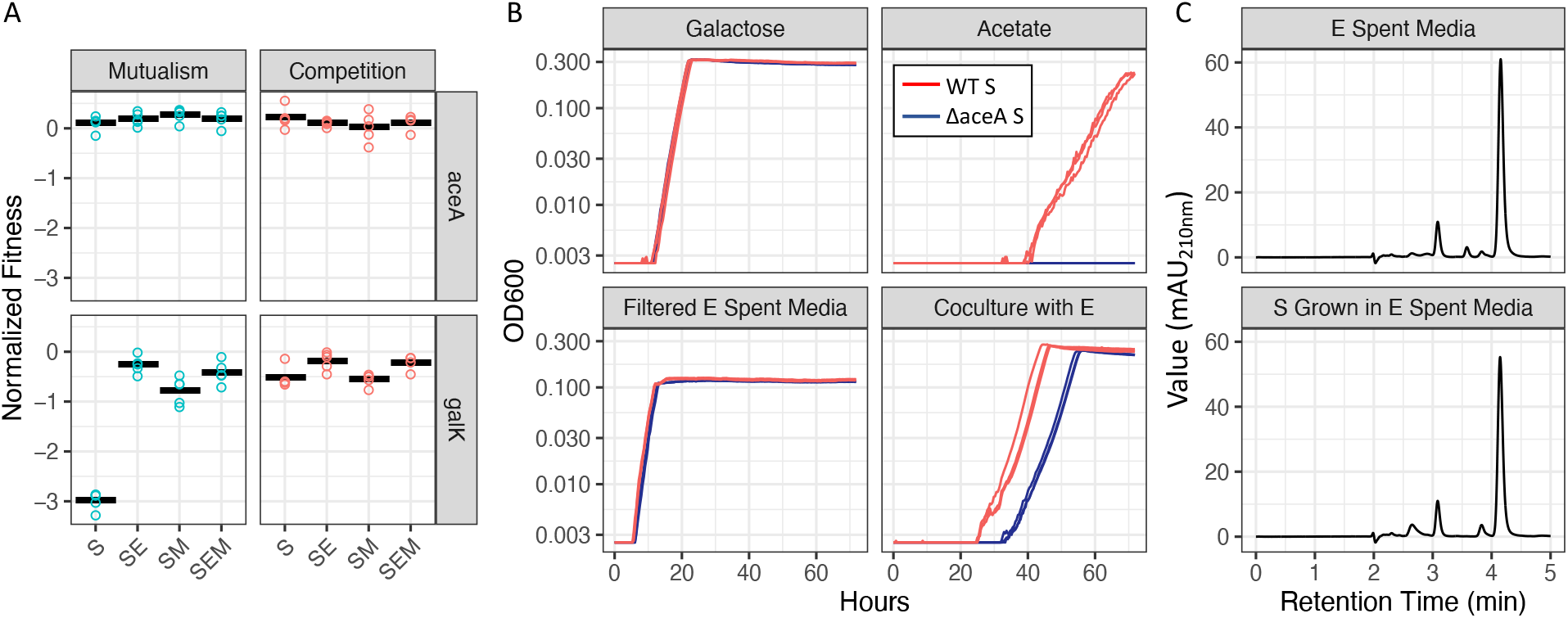
Unexpected carbon catabolism fitness patterns. A) Normalized BarSeq fitness scores for a representative acetate utilization gene (*aceA* top) and galactose utilization gene (*galK* bottom) under mutualistic (blue) and competitive conditions (red). The horizontal line represents the mean fitness for each gene in each community. B) Growth curves (OD600) of wild type (red) and *aceA* mutant *Salmonella* (blue) grown in minimal media containing galactose (top left) and acetate (top right) as the sole carbon source. In the bottom left panel growth of the two strains in filter sterilized spent media prepared from mid log *E. coli* (lactose + methionine minimal media). In the bottom right panel, growth in mutualistic co-culture with *E. coli* in lactose minimal media. C) HPLC chromatograms for *E. coli* spent media (top) and *S*. *enterica* grown on *E. coli* spent media (bottom). Standards were used to optimize this method to measure acetate, butyrate, citrate, formate, lactate, propionate, pyruvate, and succinate.

We further tested the importance of acetate in the cross-feeding interaction by generating a knockout in *S. enterica* to block acetate consumption. As expected, knocking out *aceA* prevented growth on acetate but had no effect on growth on galactose (Fig. 6B). An *E. coli* co-culture with the Δ*aceA* had 10% less final biomass (t-test, p < 0.01) and the lag time was longer compared to the co-culture with WT *S. enterica* (Fig. 6B). Similarly, on *E. coli* spent media, we found that the Δ*aceA* grew slightly (but significantly) slower (t-test, p < 0.001) and to a lower final OD600 (t-test, p < 0.05) than the WT *S. enterica* (Fig. 6B). Consistent with these findings, HPLC analysis found very little change in acetate or other organic acids when *S. enterica* was grown in the *E. coli* spent media (Fig. 6C).

### Vitamin and amino acid metabolism

Loss of biosynthetic genes for vitamins (B1, B5, B6), isoleucine and the co-factor NAD were less deleterious in mutualistic co-culture than monoculture on galactose (Fig. 7A). These results suggest that partner species may provide these nutrients to *S. enterica.* However, in our competitive environment when *S. enterica* was grown in monoculture on succinate we also saw that losing these genes had little impact on fitness. To further investigate these effects *in vitro*, we created new knockouts in *S. enterica* for critical genes in isoleucine (*ilvA*) and vitamin B5 (*panC*) biosynthesis.

**Figure 7.**
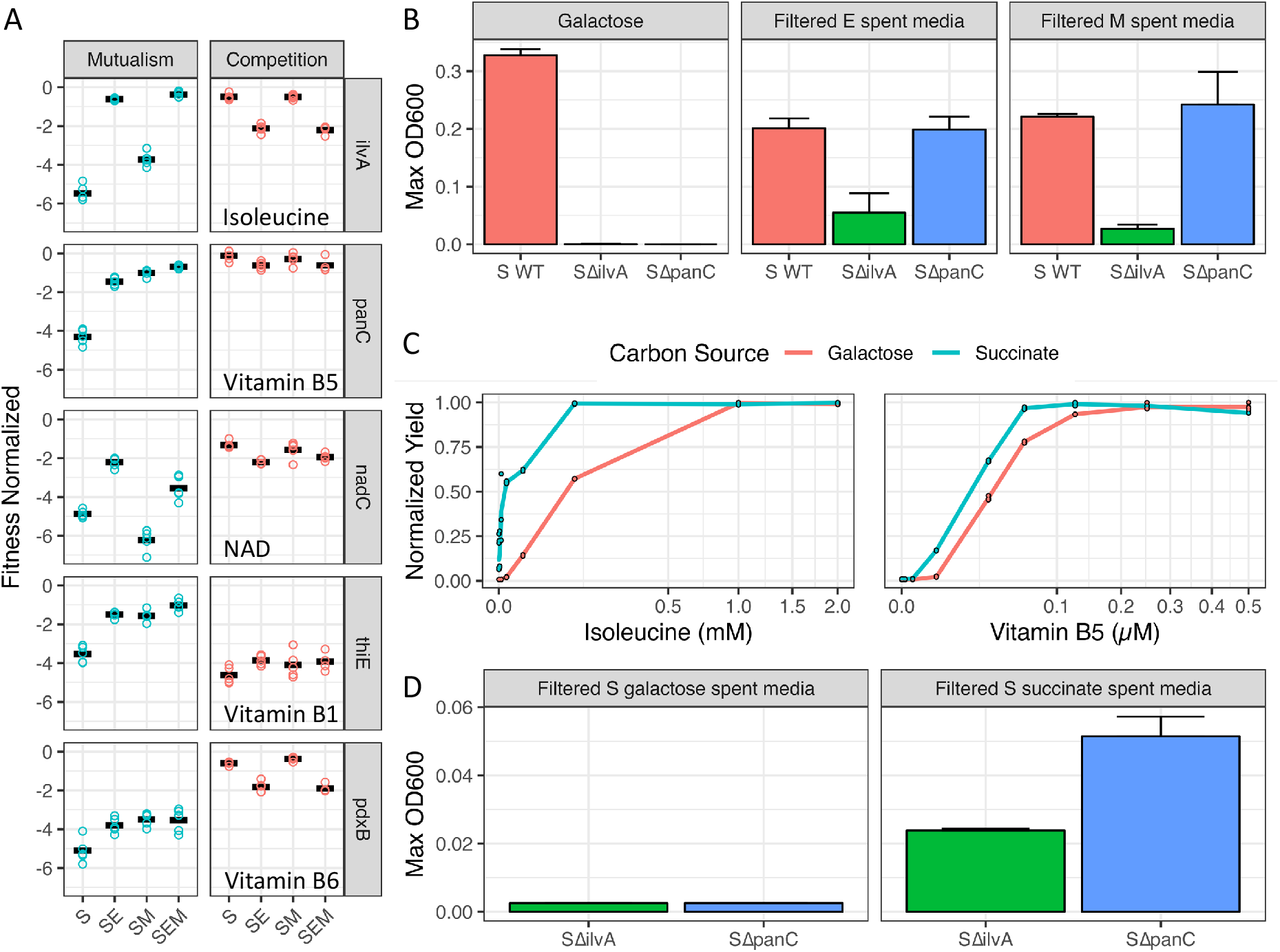
Effect of ecology on vitamin, cofactor, and amino acid metabolism gene fitness. A) Normalized gene fitness scores for representative genes involved in biosynthesis. B) Final OD yield of wild type and mutant *S. enterica* (S) on fresh galactose media (left), *E. coli* (E) spent media (middle), and *M. extorquens* (M) spent media (right). C) *S. enterica ilvA* mutant (left) and *panC* mutant (right) yield on galactose and succinate minimal media with varying amounts of supplementation. Note that the y-axis is the yield normalized to the maximum yield for each carbon sources (yield at concentration/maximum yield) and the x-axis is square root transformed. All experiments were performed in triplicate. D) Final OD yield of mutants Δ*ilvA* and Δ*panC S. enterica* grown in spent media prepared from wild type *S. enterica* grown to mid-log in galactose (left) or succinate (right) minimal media.

Knockout constructs supported that mutualism altered the need for isoleucine and vitamin B5 biosynthesis. Neither Δ*panC* nor Δ*ilvA* could grow in isolation in galactose minimal medium (Fig. 7B); however they could be rescued by either B5 or isoleucine (Supp. Fig. 4). Additionally, each mutant could be at least partially rescued by growth in spent media from *E. coli* or *M. extorquens* (Fig. 7B right). These data are consistent with *S. enterica* acquiring multiple metabolites from each mutualistic partner.

We next investigated the neutrality of Δ*panC* and Δ*ilvA* mutants in our competitive monoculture treatment. We found that the Δ*panC* mutant was unable to grow in succinate. Interestingly, on succinate, the Δ*ilvA* mutant was only able to grow after ∼60 hours. Literature suggests that this growth is due to the gene *tdcB* (28), encoding an enzyme that yields 2-oxobutanoate (similar to *ilvA*) but only in the absence of sugar and oxygen. That said, over a 48-hour period, we found that neither mutant was capable of substantial growth on succinate. Furthermore, the mutants needed more isoleucine or vitamin B5 to maintain growth on galactose than on succinate (Fig. 7C). We also found that both mutants grew to higher yields in spent media from wildtype *S. enterica* grown on succinate than on galactose (Fig. 7D). These results suggest that the mutants could acquire needed metabolites from wildtype *S. enterica* (as well as from mutualistic partners), but that both the demand and secretion of the metabolites changed as a function of carbon source.

## Discussion

Using a transposon library in a defined microbial consortium, we found that ecological interactions have distinct effects on the impact of mutations in *S. enterica.* The average fitness of all mutants was closer to neutral when *S. enterica* was engaged in mutualism than when it was grown alone. However, this buffering was not observed when *S. enterica* was in competition with the same species. We also found that the impact of mutations in a three-species community could be well predicted from the average impact of the mutations in each two-species co-culture. Investigation of the impact of specific gene disruptions in the different ecological communities illuminated the mechanisms of species interactions. As expected, nitrogen uptake genes were critical when *S. enterica* was reliant on ammonium secreted by *M. extorquens.* In contrast, the fitness effect of gene disruptions did not support the expected mechanisms of carbon exchange between *E. coli* and *S. enterica.* Finally, the transposon mutant fitness data highlighted that additional essential metabolites can be obtained from other cells, but both the supply and demand of these metabolites was impacted by the carbon environment.

Mutualism buffered the effect of mutations in our system. The mean fitness of mutants was higher in mutualism than monoculture for all conditions despite the fact that the partner species and mechanisms of mutualism were distinct. The buffering of fitness effects in co-culture could be driven by generic effects on growth rate, as *S. enterica* grows more slowly in all mutualisms than it does in monoculture (Supp. Fig. 2). However, we also demonstrated that both *E. coli* and *M. extorquens* rescue the growth of specific mutants through the release of metabolites.

Mutualism and competition had distinct impacts on the average effect of mutations. Competition did not alter the average effect of mutations relative to monoculture. The lack of impact on average fitness could in part be explained by the strength of competition. In particular, *M. extorquens* is a relatively weak competitor, though it does significantly reduce the total biomass of *S. enterica* relative to monoculture (t-test, p < 0.05). Independent of the strength of interactions, the competition results highlight that the ability of mutants to obtain essential metabolites from other species is strongly influenced by how species are interacting. One cannot predict the fitness effects of losing biosynthetic pathways simply from species presence in a community – instead, it is critical to know how species are interacting.

Our results suggest that the impact of mutations in complex communities can be predicted from the impact of mutations in simpler co-cultures. The fitness effect of mutations in the 3-species community could be well predicted from fitness effects in each 2-species co-culture. Our results provide some contrast to findings with an *E. coli* BarSeq library in co-culture with cheese rind microbes. Morin *et al.* found that the impact of a species on the fitness of a mutant in a focal species was influenced by the presence of a third species (i.e., higher order interactions (15). Consistent with Morin *et al.* we found that the effects of species on mutant fitness were often not additive. However, despite the frequent subadditivity of species effects in our system, an additive model was still able to provide strong predictions of fitness in the 3-species community. The prediction could be improved even more by calculating the average fitness effect over the two 2-species communities. Our results suggest that even in complex communities, selection, and therefore evolutionary dynamics may be predictable with data from simpler systems.

BarSeq analysis allowed us to test expected mechanisms of species interaction in our synthetic community. The fitness of nitrogen acquisition mutants supported our expectation that *S. enterica* acquires ammonium from *M. extorquens* in the mutualism. We were encouraged our fitness results mirrored findings from other systems in which *E. coli* acquires ammonium from partners (12).The carbon exchange between *E. coli* and *S. enterica* remains less clear. In contrast to our expectations, the BarSeq results suggest that neither acetate nor galactose are the primary sources of carbon provisioned by *E. coli*. Furthermore, spent media analysis suggests that other organic acids are not the primary source of carbon either. We now suspect that a sugar other than galactose is likely exchanged; however, further work will be required to identify the source(s).

We also identified unknown metabolite exchange in our mutualism. Disruption of genes involved in vitamin, amino acid, and cofactor biosynthesis was less deleterious in mutualism than in monoculture. To better understand how mutualistic interactions improved fitness, we generated knockouts for two genes that had large fitness improvements in mutualism: *ilvA* – a gene involved in isoleucine biosynthesis, and *panC –* a gene involved in the production of vitamin B5 (pantothenate). While neither mutant grew in isolation on galactose minimal media, each grew in the presence of either *E. coli* or *M. extorquens.* This suggests that *E. coli* and *M. extorquens* excrete isoleucine and vitamin B5 (or precursors for these molecules). Several previous studies have documented rescue of auxotrophs in co-culture. In *E. coli* it has been shown that amino acids (29,30), cofactors, and vitamins (31) can be obtained from other cells. Additionally, the phototrophic bacterium *Rhodopseudomonas palustris* has been shown to rescue *E. coli* with knockouts of purine, vitamin B6, and NAD biosynthesis (12). Gude *et al.* suggested that overproduction of essential nutrients may be favored to avoid bottlenecks in metabolism (32). In our system *E. coli* and *M. extorquens* each rescued a similar set of *S. enterica* auxotrophs supporting that there are a set of metabolites commonly released into environments by different taxa.

Carbon sources in our media impacted the ability of auxotrophs to be rescued by other cells. Many biosynthetic genes that were critical in monoculture on galactose were far less important when *S. enterica* was grown in monoculture on succinate. This led us to the hypothesis that mutants could obtain essential metabolites from other *S. enterica* cells, but that the carbon source changed the amount of metabolites released or the amount of metabolites required to rescue growth. Experiments with Δ*panC* and Δ*ilvA* mutants supported that growth on succinate led to both more excretion of essential metabolites by *S. enterica* and less metabolite being required for mutant growth. These results highlight that carbon source strongly impacts cross-feeding of nutrients, and the degree to which cross-feeding buffers the impact of gene loss.

There are a number of caveats when considering the general applicability of our results regarding mutualism, competition, and the distribution of fitness effects. First, we only studied one form of mutation: gene disruption. We did not study basepair changes which may be more common drivers of adaptation. Though it is worth noting that a recent study demonstrated that distributions of fitness effects derived from transposon mutagenesis were predictive of evolutionary dynamics observed in *E. coli* (7). Second, the carbon source varied across our treatments, and we demonstrated that carbon source was sufficient to alter the fitness effects of mutants. Encouragingly, we found that mutualism had consistent impacts of reducing the average effect of mutations even though the carbon source changed between mutualistic treatments from galactose to the carbon that *S. enterica* obtains from *E. coli*.

The challenges of predicting evolution in communities has long been appreciated (33). However, we observed that the fitness effects of mutations could be predicted to a degree in synthetic communities. All mutualisms we evaluated buffered the impact of gene loss, and indeed there was substantial overlap in the functions that were buffered by different mutualistic species. Furthermore, fitness in co-cultures was sufficient to predict fitness in more complex microbial communities. Our results suggest that synthetic communities of mutualists may often be more robust to genetic perturbations than synthetic communities of competitors. Additionally, our results suggest that study of pairwise interactions may ultimately allow us to understand, predict and ultimately manage evolutionary dynamics even in complex natural communities.

## Acknowledgements

Acknowledgements

The authors thank A. Bisesi, X. Xiong, and V.G. Martinson for helpful comments on the manuscript. The authors would also like to thank the Deutschbauer lab for the generous gift of the pKMW3 donor library and providing answers to questions about BarSeq. Finally, the authors would like to acknowledge BEI resources for providing 10428S *S. enterica* mutant strains. This work was supported by the National Institutes of Health R01-GM121498 to W.R.H.

---

**Supplemental Figure 1.**
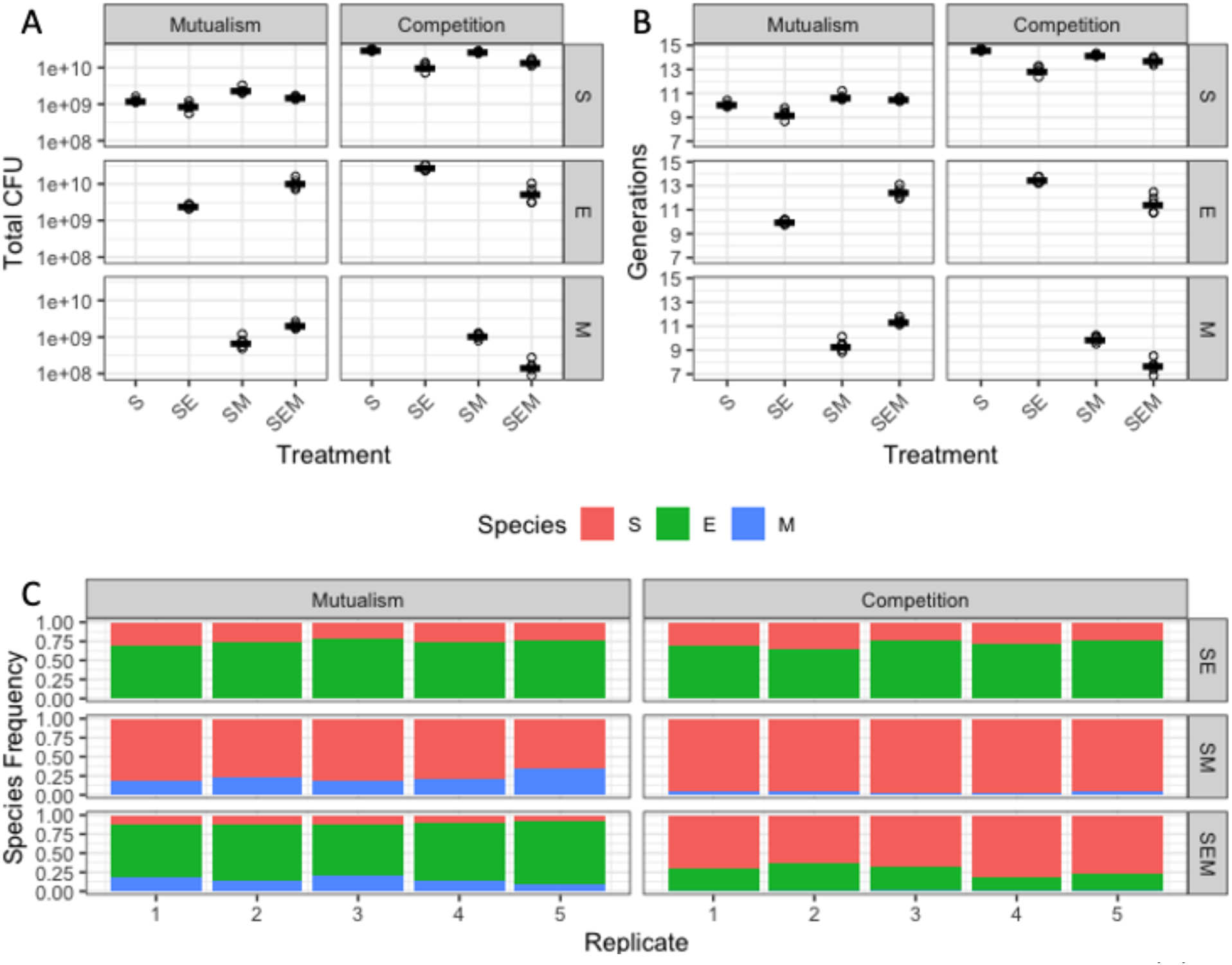
Species Population Sizes and Generations. Total population size (A) and number of generations (B) for each species (indicated by y-axis faceting) in each treatment. C) Species frequency for each co-culture replicate.

**Supplemental Figure 2.**
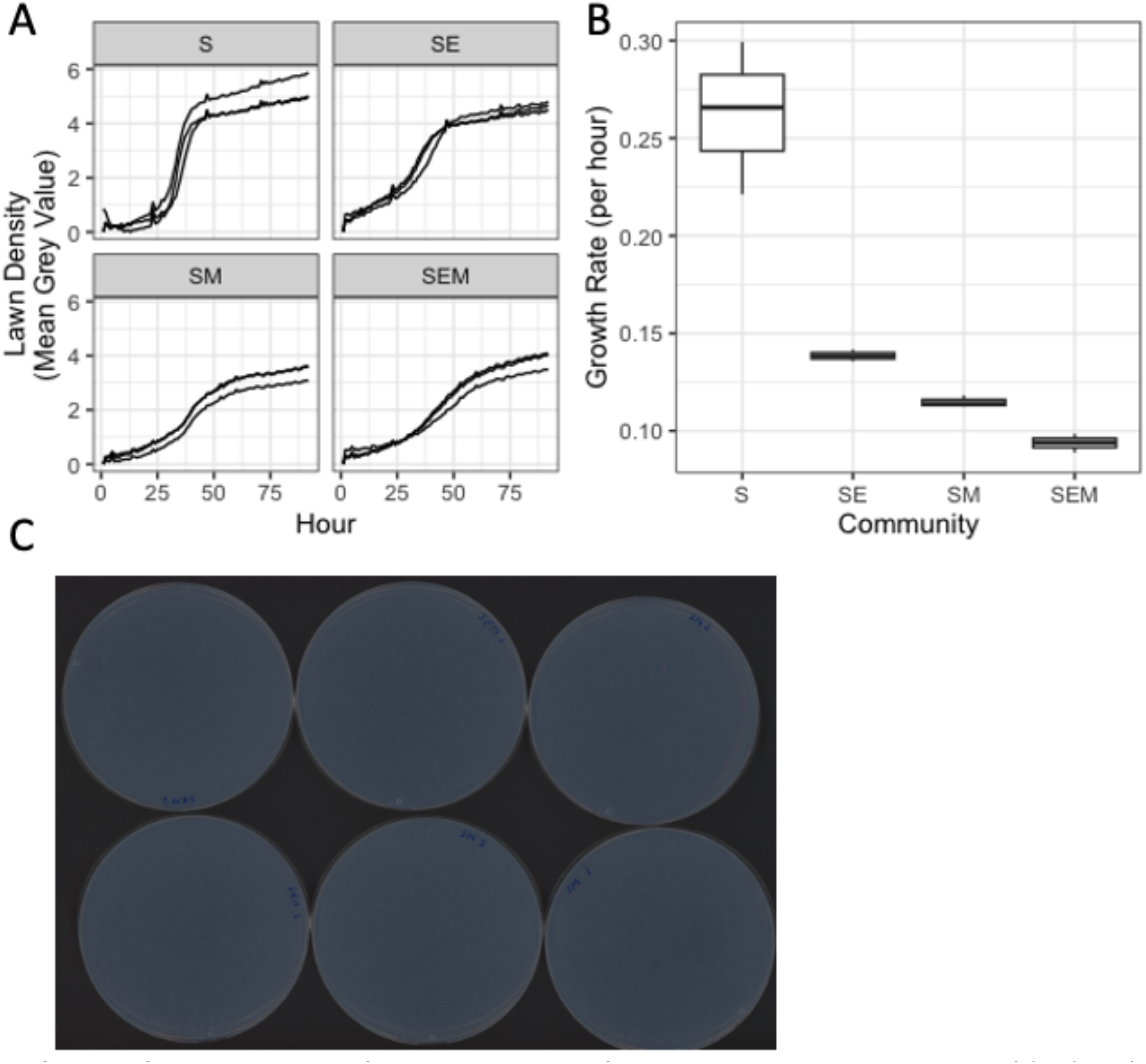
Growth on agar. Grow of communities on agar measured by hourly automated scanning of each mutualistic community in triplicate. Density of bacterial lawns was approximated by converting each plate image to greyscale, then calculating the mean grey value through time (A). Growth rates were fit using baranyi curves (B) – S monoculture’s growth rate is significantly greater than mutualistic co-culture (Tukey HSD, p < 0.05 for S vs SE, S vs SM, S vs SEM). C) Example image of petri dishes on scanner.

**Supplemental Figure 3.**
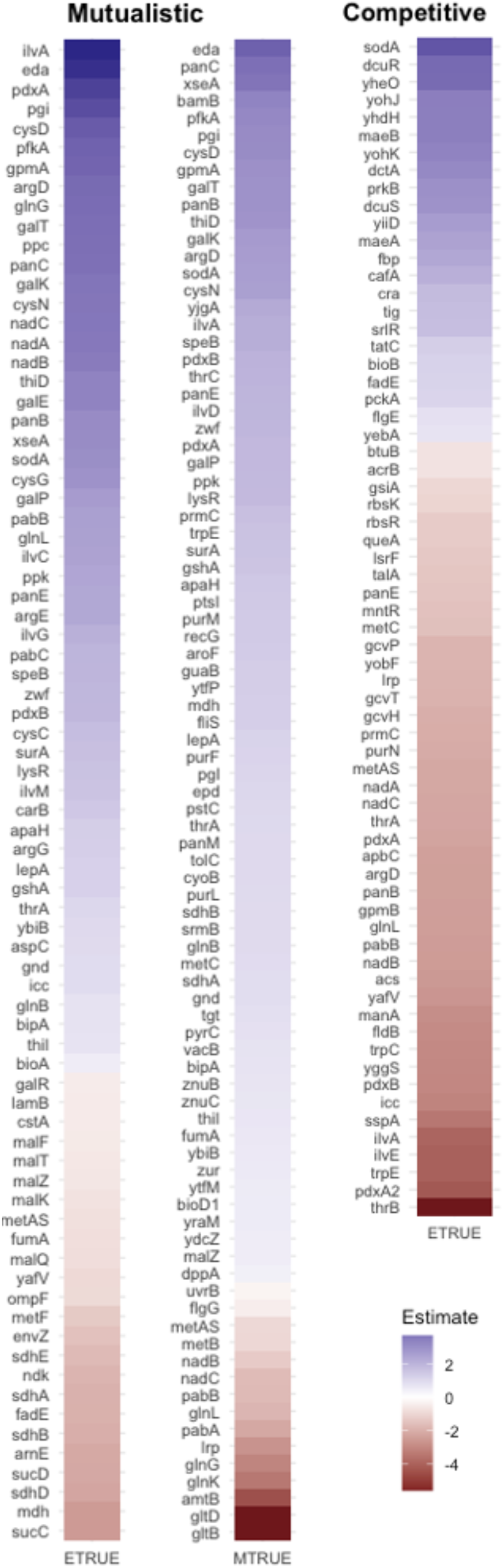
Genes with Significant Fitness Effects Heatmap. Genes with significant effects (adj. p-val < 0.05) in the presence of E or M in mutualistic (left) and Competitive (right) conditions. Color indicates the linear regression estimate – fitness improvements (relative to monoculture) are indicated in blue, while decreased fitnesses are indicated in red. Note that no genes were significant for the competition in the presence of M.

**Supplemental Figure 4.**
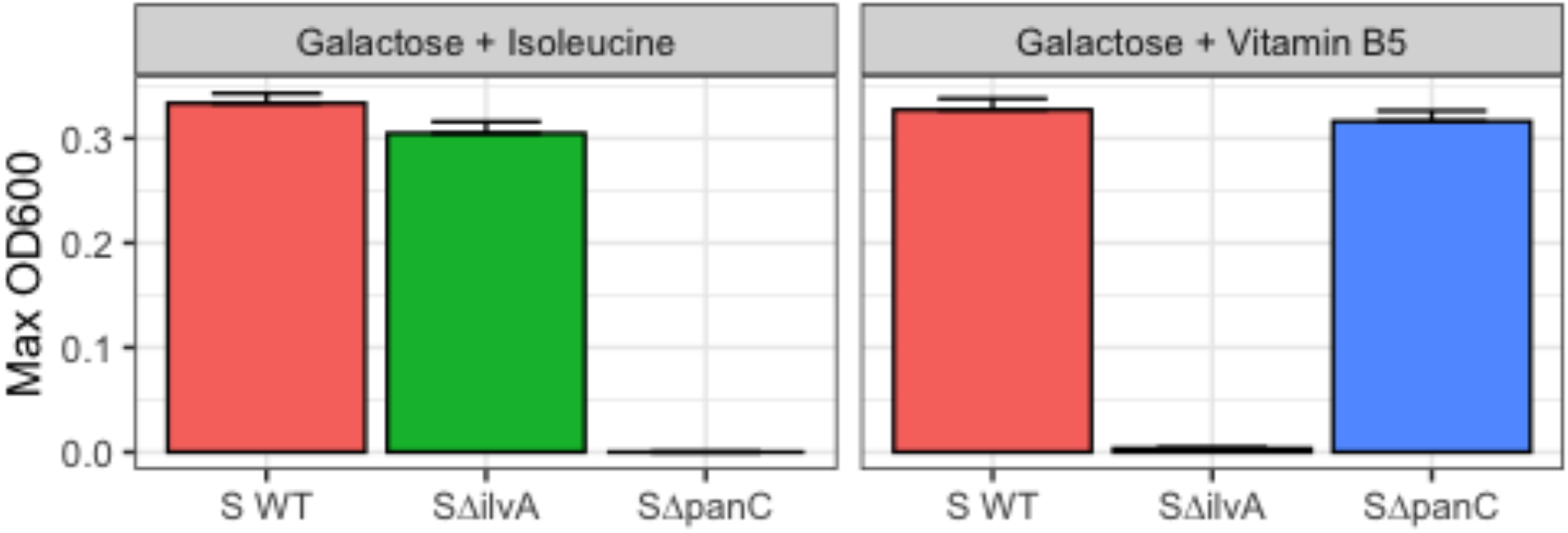
Amino Acid and Vitamin Supplementation Restores Auxotroph Growth. End yields of Wild type and Δ*ilvA* / Δ*panC S. enterica* grown in galactose Hypho broth supplemented with 1 mM isoleucine (left) and 0.5 µM Vitamin B5 (right). Each treatment combination was performed in triplicate.

